# Functional characterization of splicing regulatory elements

**DOI:** 10.1101/2021.05.14.444228

**Authors:** SI Adamson, L Zhan, BR Graveley

## Abstract

**Background:** RNA binding protein-RNA interactions mediate a variety of processes including pre-mRNA splicing, translation, decay, polyadenylation and many others. Previous high-throughput studies have characterized general sequence features associated with increased and decreased splicing of certain exons, but these studies are limited by not knowing the mechanisms, and in particular, the mediating RNA binding proteins, underlying these associations.

**Results:** Here we utilize ENCODE data from diverse data modalities to identify functional splicing regulatory elements and their associated RNA binding proteins. We identify features which make splicing events more sensitive to depletion of RNA binding proteins, as well as which RNA binding proteins act as splicing regulators sensitive to depletion. To analyze the sequence determinants underlying RBP-RNA interactions impacting splicing, we assay tens of thousands of sequence variants in a high-throughput splicing reporter called Vex-seq and confirm a small subset in their endogenous loci using CRISPR base editors. Finally, we leverage other large transcriptomic datasets to confirm the importance of RNA binding proteins which we designed experiments around and identify additional RBPs which may act as additional splicing regulators of the exons studied.

**Conclusions:** This study identifies sequence and other features underlying splicing regulation mediated specific RNA binding proteins, as well as validates and identifies other potentially important regulators of splicing in other large transcriptomic datasets.

## Background

Alternative pre-mRNA splicing is a key process which can regulate gene product abundance and increase proteome diversity. While significant progress has been made in understanding the mechanistic underpinnings of splicing regulation, we still lack a complete understanding of how sequence variants and RBP dysregulation can impact splicing. Certain sequence features such as 5’ and 3’ splice sites are relatively well understood, and in particular, the importance of the 5’ and 3’ splice site dinucleotides required for most pre-mRNA splicing reactions. Most variant impact prediction pipelines include prioritization of variants impacting these dinucleotides, and some include prioritization of variants proximal to these splice sites outside of splice site dinucleotides [1–4]. Variants outside of these sequences however have been shown to impact splicing and can be underpinnings of monogenic and polygenic diseases [5–7]. In the context of some Mendelian diseases, like spinal muscular atrophy, progress has been made in understanding how specific RBPs can regulate disease underlying splicing events and how variants can disrupt these interactions [8], however these tend to be exceptions. Broadly, understanding how variants can disrupt these RBP-RNA interactions which regulate splicing, or splicing regulatory elements (SREs), has remained understudied. Understanding the how variants disrupt SREs and the downstream effect on splicing is an important step to understanding how novel or unstudied variants may be related to phenotypes.

Two main approaches have been employed to understand the impact of genetic variants on splicing regulation in a high throughput manner, these include machine learning approaches and massively parallel splicing reporter assays (MPRAs). The application of machine learning approaches to understanding the sequence features underlying splicing regulation (also referred to as the “splicing code”) has become a fruitful endeavor in recent years, which may reflect the rapid development and accessibility of deep-learning and neural networks more broadly. These approaches have been used to predict the impact of genetic variants on splicing including alternative splicing, and the use of novel splice sites [9–14]. These approaches are useful and reasonably accurate, however predictions are not always reliable. MPRAs which have been employed to study the impact of variants on splicing employ diverse approaches including the use of random sequences, and encoding genetic variants observed in the general population as well as in the context of disease [15–23]. MPRAs, while having their caveats, are able to test the impact of genetic variants on splicing directly. However these studies have mostly examined the impact of RBPs from the level of *in vitro* motifs or broad sequence features which are RBP agnostic. Knowing which RBPs likely mediate variant-induced splicing changes is important to develop interventions to change splicing to a wild-type state in the context of disease, and to understand more broadly the underpinnings of splicing regulation.

In this study we sought to understand the role of RBPs in splicing regulation, and in particular, how sequence variants impact pre-mRNA splicing. To do this, we first explored the ENCODE RNA binding protein datasets, which consists of RBP binding data (eCLIP), RBP knockdown RNA-seq, and RNA bind-n-seq [24–26]. This is a source of rich datasets which when integrated, can identify RBP-specific SREs and attempt to refine RBP binding sites using motif information. Here we use this data in order to identify functional SREs and features which underlie RBP knockdown-induced splicing changes. We next surveyed the underlying sequence requirements of these SREs on splicing regulation using a massively parallel splicing reporter assay we previously developed called Vex-seq [15] and validated a small subset of these results using base-editing SRE in the endogenous locus. Finally, we validate the role of ENCODE identified RBPs and the potential regulatory role of other RBPs on the exons we studied through examining TCGA and GTEx. Together, these results serve to validate many of the SREs identified by ENCODE and set forth a strategy for globally surveying SREs on a genome-wide scale.

## Results

### Identification and characterization of splicing regulatory elements in ENCODE datasets

To identify RBPs that behave as splicing regulators in ENCODE datasets, we integrated eCLIP and RBP knockdown RNA-seq datasets to understand the relationship between direct RBP binding and splicing regulation. We analyzed these paired datasets primarily for annotated splicing regulatory proteins and novel RBPs (based on annotations in [26]), which include 51 proteins in K562 cells and 44 proteins in HepG2 cells (datasets listed in Additional File 1). RBPs with a direct impact on splicing regulation would likely show an enrichment of binding sites near regions with splicing changes upon RBP knockdown relative to other binding sites throughout the transcriptome. To quantify this, we used Fisher’s exact test to identify enrichment of binding within 300 bases of exons changing after knockdown compared to binding near exons which did not change splicing with RBP knockdown. Certain members of well characterized splicing regulatory protein families, like HNRNPs and SR proteins, tended to cluster together and showed similar directionality upon knockdown (Figure 1a). SR proteins tended to show the most significance when binding to exons which decreased splicing upon RBP knockdown, and the opposite was generally true for HNRNPs. 84.2% of RBPs with annotated roles in splicing regulation, tending to have enriched binding near exons which change splicing with RBP knockdown (Bonferroni-corrected p ≤ 0.01 in at least one region), while only 42.1% of RBPs without known roles in splicing regulation showed this trend. UCHL5 and FXR1 were notable RBPs without known splicing regulatory roles which showed significant dependence between binding and splicing regulation. While some known splicing regulatory proteins, like TRA2A, didn’t show any enrichment of binding at sites of changed splicing.

**Figure 1).**
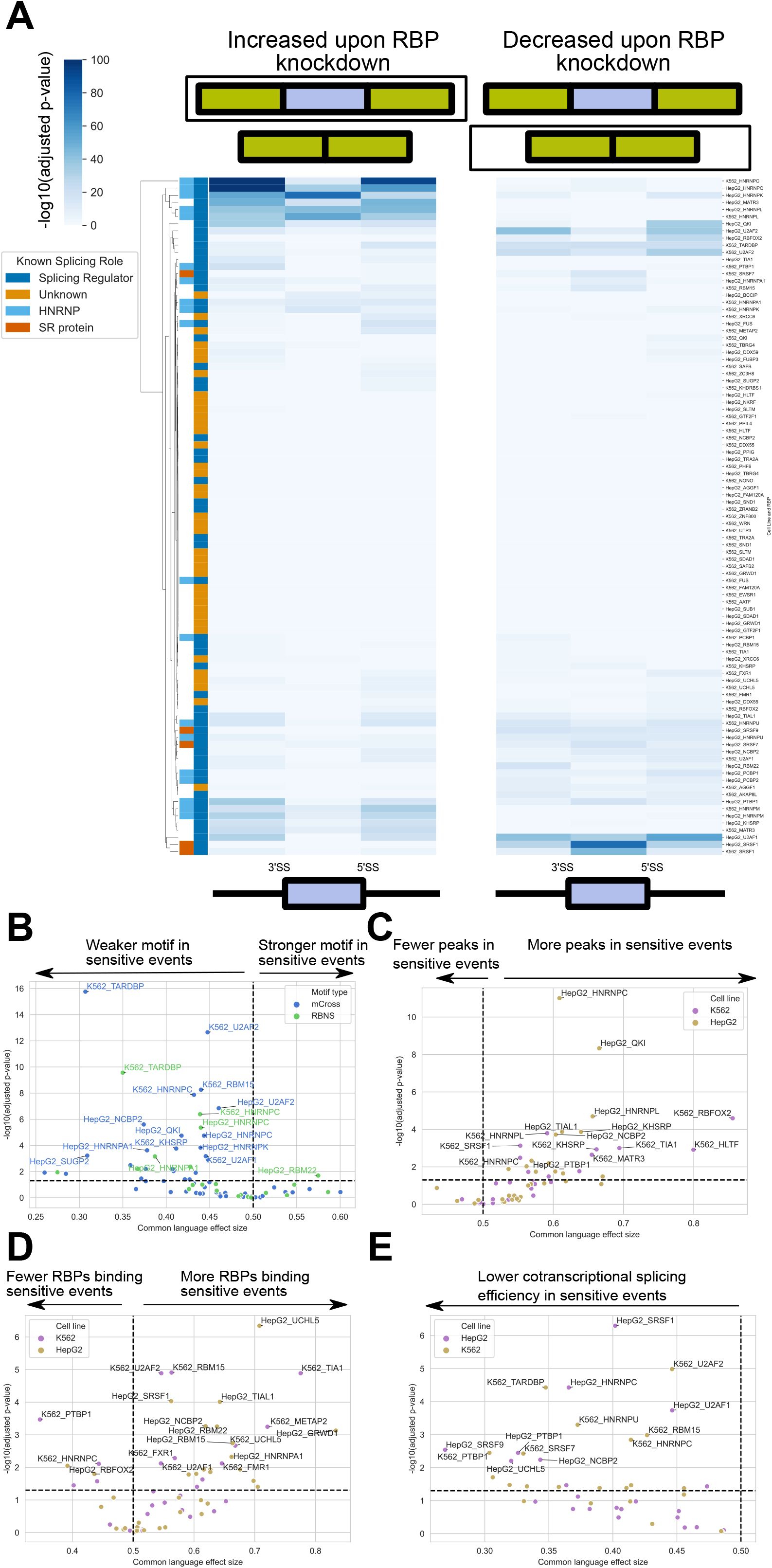
Characterization of RBP-specific splicing regulatory elements in ENCODE data. **a)** Clustered heatmap showing the significance of dependence between binding and splicing knockdown sensitivity assessed by Fisher’s exact test. -log_10_(Bonferroni corrected p-values) in each region for upregulated upon knockdown (left) and downregulated upon knockdown (right) is the quantity being clustered. Known splicing regulatory proteins are annotated in dark blue, while proteins with no known role in splicing regulation are in orange. HNRNPs are highlighted in light blue, while SR proteins are highlighted in red. Volcano plots showing the significance and common language effect size for differences in **b)** motif strength, **c)** number of peaks from the designated RBP, **d)** number of other RBPs binding to the splicing event, and **e)** co-transcriptional splicing efficiency between robust and sensitive splicing events. Significances and effect size are assessed by Mann-Whitney-U test and Benjamini-Hochberg correction.

Understanding which features make certain RBP-bound exons change splicing upon knockdown is important to understand the underpinnings of splicing regulation. We will refer to RBP-bound exons which change splicing with knockdown of the same RBP as “sensitive” and RBP-bound exons which do not change splicing as “robust”. There are a number of possible reasons which could underlie splicing events showing sensitivity to RBP knockdown, including the concentration of RBP in the cell, the binding affinity of the RBP to its RNA substrate, the temporal regulation of splicing, and the presence of other splicing regulatory RBPs binding nearby. We investigated a number of these features and their role in differentiating sensitive and robust splicing events.

We hypothesized that splicing events that are robust to knockdown have a higher binding affinity of the RBP to their cognate SREs than those which are sensitive to knockdown. To study this, we examined the motif strength of peaks near sensitive and robust events by examining mCross motif position specific scoring matrices (PSSMs) motif scores and RNA bind-n-seq (RBNS) derived RBPamp affinities in 30 base windows around the peaks [24, 27]. mCross [27] is a database of RBP motifs derived from eCLIP data, and RBPamp (Jens M. et al., submitted) is a method which can estimate dissociation constants of RNA-RBP interactions through computational modeling of RBNS data. Consistent with our hypothesis, more RBPs binding near sensitive events show weaker motifs/affinities compared to robust peaks compared to the opposite behavior (Figure 1b). While motif strength may indicate whether an RBP will interact with a particular motif, multiple binding sites for a single RBP can be observed regulating a single splicing event. To explore whether additional binding sites of a single RBP are important for determining whether a splicing event is sensitive to knockdown, we examined how many peaks for a given RBP were in sensitive and robust events for particular datasets (Figure 1c). Many datasets tended to show a bias for having more binding sites in splicing events sensitive to knockdown when compared to those which were not. Additional layers of RBP splicing regulation may underlie whether a splicing event changes with RBP knockdown. To examine the role of other RBPs, we asked whether for each dataset, if sensitive events were regulated by more or fewer RBPs compared to robust events (Figure 1d). Many datasets with meaningful trends showed that there were more RBPs binding to sensitive events, relative to robust ones, however some notable splicing regulators, like PTBP1, HNRNPC, and RBFOX2 showed the opposite trend.

Another possible difference between sensitive and robust events could relate to the timing of splicing relative to transcription. Previous studies have observed that the removal of introns flanking constitutive exons is typically more efficient and faster than the removal of introns flanking alternative exons [28, 29]. It has also been shown that elongation speed of RNA Pol II can regulate alternative splicing [30–32]. This led us to hypothesize that the kinetics of intron removal may be a factor influencing whether splicing events are robust or not to RBP knockdown. Co-transcriptional splicing efficiency for introns flanking alternative exons were obtained from [32] and compared between sensitive and robust events. We observe that in many RBP datasets, there is a significantly lower co-transcriptional splicing efficiency in sensitive events compared to robust ones (Figure 1e).

### Characterization of the functional impact of mutations on empirically identified splicing regulatory elements

Characterizing the sequence determinants of functional SREs is key to predicting the impacts of genetic variation on splicing and understanding the underlying molecular processes. To this end, we utilized a slightly modified version (see Methods) of a high throughput splicing reporter assay we previously established, called <u>V</u>ariant <u>ex</u>on <u>seq</u>uencing (Vex-seq), to study the sequence determinants of SREs in a massively parallel manner (Figure 2a). We overlapped SREs which were sensitive to RBP knockdown with exons which could be studied in Vex-seq due to the size constraints of pooled synthetic oligos, and which had the RBP binding sites of the SRE in a testable window. The distribution of testable exons, given the size constraints, generally reflected the distribution of all sensitive exons for each dataset (Supplemental Figure 1). After calling eCLIP peaks using multiple pipelines, we used de-novo motifs identified from the eCLIP peaks to attempt to refine the location of the actual binding sites. We then designed di-nucleotide mutants around and including the putative binding sites for the RBPs within the SREs. In total, 33,317 unique sequences were designed across 259 unique exons to study 355 distinct SREs (Additional file 2). The SREs selected for these studies are recognized by well-known splicing regulators, as well as some with less characterized roles in splicing regulation (Figure 2b). These sequences were assembled in parallel into a library of Vex-seq reporter plasmids as previously described with minor modifications [15] (see Methods for details). Sequences were designed with three barcode replicates. Analysis of the representation of each of the barcodes in the final plasmid library pool showed that the top decile of barcodes was represented 7.75 times as much as the bottom decile (also known as skew ratio) (Supplemental Figure 2a). 825 barcodes dropped out during assembly of the plasmid pool, however these only represent 0.825% of all barcodes. To ensure the sequences were reflective of the sequences designed, we called variants on each barcode and filtered out barcodes with variants represented in more than 20% of the reads associated with each barcode (Supplemental Figure 2b). The Pearson correlation of Percent Spliced In (Ψ) and change in Ψ between variant and reference (ΔΨ) between biological replicates of the same cell line was ≥ 0.95 and ≥ 0.73, respectively (Supplemental Figure 2c&d).

**Figure 2).**
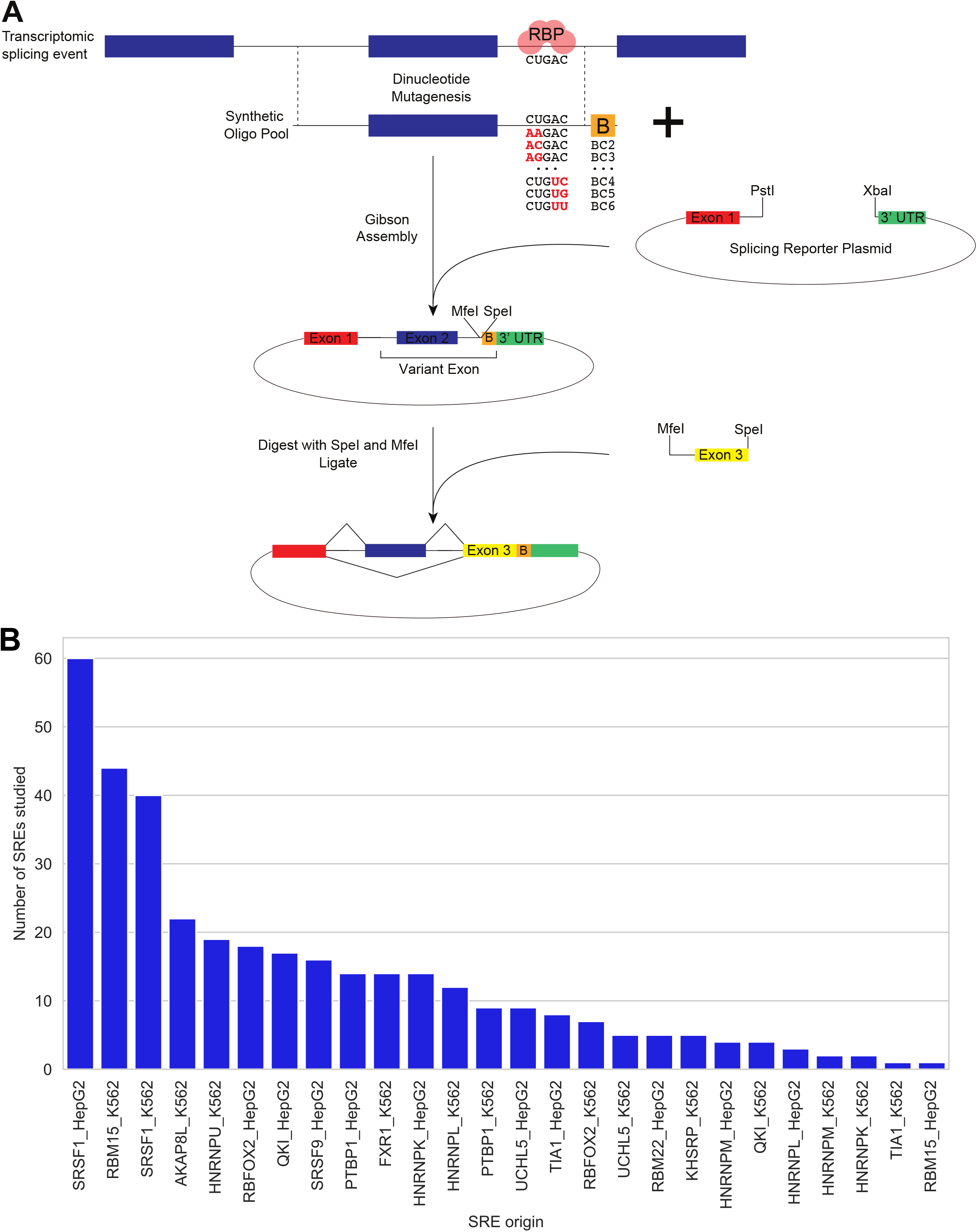
Experimental design of Vex-seq to characterize the sequence determinants of SREs. **a)** An illustration showing a simplified experimental schematic of Vex-seq and its application to study SREs. **b)** A histogram of SREs and their dataset origins which were designed to be assayed using Vex-seq.

Most designed variants tended not to disrupt splicing, however high magnitude ΔΨ outliers were observed in many locations with 5.9%, 5.44%, 4.27% of variants with |ΔΨ| > 20 in upstream introns (> 20 bases upstream of 3’ splice sites), exons, and downstream introns (> 20 bases downstream of 5’ splice sites), respectively (Figure 3a). Some designed variants resulted in unexpected splicing outcomes including alternative 3’ splice site (A3SS) usage, alternative 5’ splice site (A5SS) usage, and sometimes an entirely unexpected exon (Figure 3b). To identify these, we used stringtie [33] to annotate novel isoforms and SUPPA2 [34] to assess the significance of the splicing change (Additional Files 3 and 4 for HepG2 and K562 respectively). Unannotated exons were detected in 73 reference sequences, however only seven had an unannotated exon spliced in more frequently than the annotated exon in the HepG2 data and eight in K562 (Supplemental Figure 3 a&b). For all significantly changing skipped exon events, (BH corrected p-value ≤ 0.05 assessed by SUPPA2) the majority of them had no direct impact on splice site strength (Figure 3c&d). This is unsurprising for annotated exons considering there were few designed variants around the 3’ and 5’ splice sites relative to other areas (see upper histogram, Figure 3a). For significantly changing A3SS and A5SS splicing events, we wouldn’t expect the same distribution of variants relative to the splice sites, considering that none of these events were specifically designed to be studied. There are considerably more A3SS events than A5SS events, this may be due to the larger window of sequences involved in 3’ splice site selection than 5’ splice site selection, or the distribution of variants around each sequence (Figure 3e&f, Supplemental Figure 4). Interestingly, designed variants which changed the 3’ splice site strength of the shorter exon (the smaller exon produced by splicing to the downstream 3’ splice site) tended to alter splicing more frequently than those which changed the 3’ splice site strength of the long exon (the larger exon produced by splicing to the upstream 3’ splice site) irrespective of whether the MaxEntScan score of the other isoform is changed (odds ratio = 2.0, p = 7.61e-15 overall; odds ratio = 2.1, p = 1.2e-07 without other splice site change by Fisher’s exact test). Additionally, 388 designed variants changed both the short and long 3’ splice site strength. Very few designed variants resulted in significant changes in the 5’ splice sites (Supplemental Figure 4).

**Figure 3).**
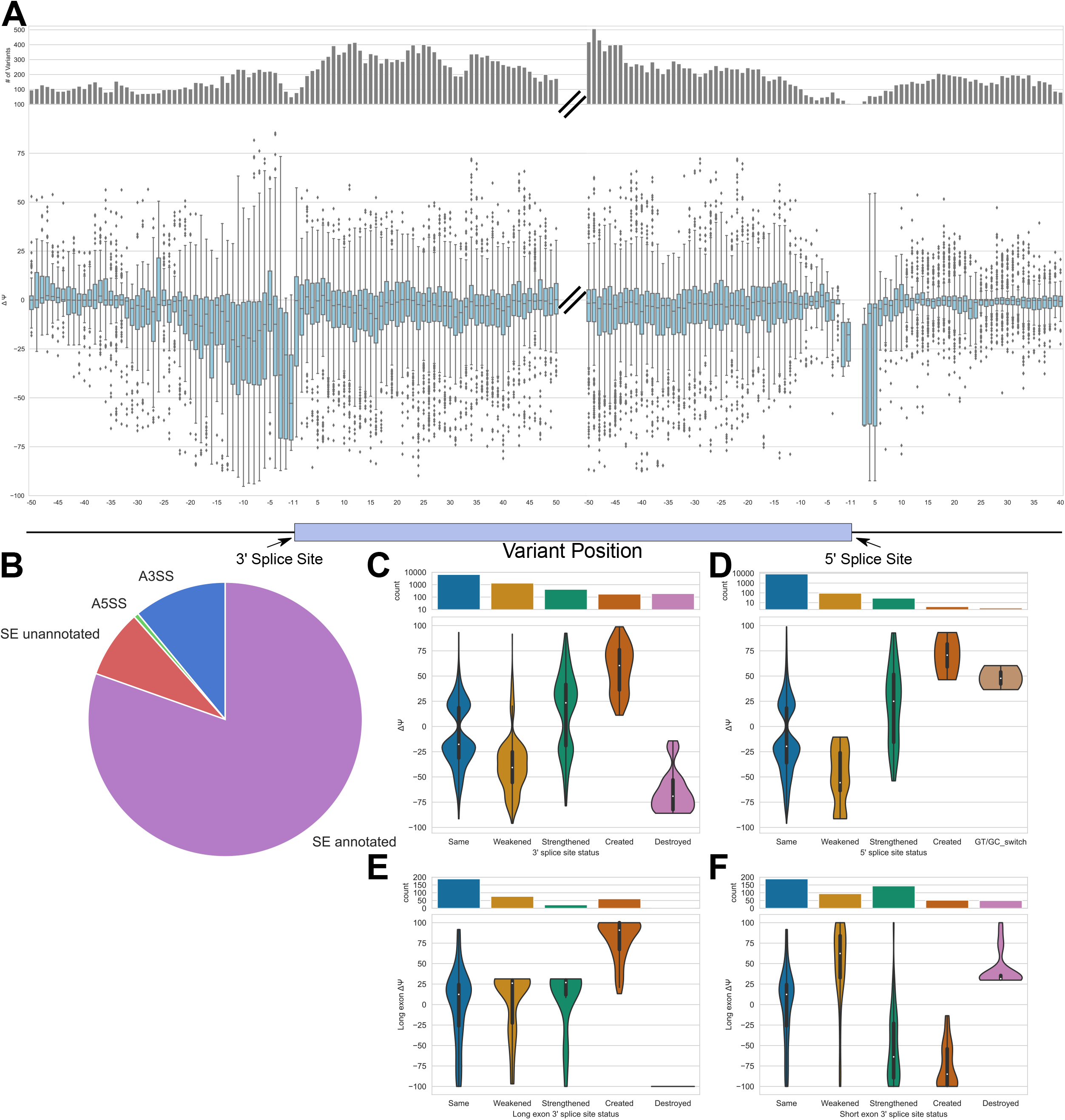
Aggregate splicing results from SRE Vex-seq data. **a)** Boxplots of ΔΨ (average of HepG2 and K562) relative to 3’ and 5’ splice sites for designed (annotated) exons. A histogram above shows the number of variants in each location. **b)** Pie chart describing the types of significantly changing splicing events (n = 9,875). Violin plots showing the distribution of ΔΨ of significantly changing splicing events assessed by SUPPA2 and broken down by whether variants changed splice site strength evaluated by MaxEntScan for **c)** 3’ splice sites in SE events, **d)** 5’ splice sites in SE events, **e)** 3’ splice sites of the long exon in A3SS events, **f)** 3’ splice sites of the short exon in A3SS events. **b-f** show data from HepG2 only. **c-f** histograms above show the number of variants in each location and only show events which do not change the other splice site strength.

To understand sequence features associated with splicing changes, we analyzed kmers and their association with ΔΨ. To this end, we analyzed pentamers and whether closely related sequences (i.e. variants designed based on the same reference sequence; ≤ 4 nucleotides different between each other) with the given kmer have a significantly different ΔΨ relative to closely related sequences without it. Pentamers were chosen because they allowed a median of 131, 50, and 36 ΔΨ measurements per kmer in exons, upstream introns, and downstream introns, respectively. A heatmap of significant kmers and their association with being spliced in or out is shown in Figure 4a (Additional file 5). 534 of all 1,024 pentamers showed a significant difference between the ΔΨ of similar sequences containing it, compared to without it. Most of these (454) were associated with changes in splicing occurring in exons, while fewer were associated with changes in downstream introns (14) and upstream introns (137). In order to compare our observations with previous datasets, we analyzed hexamers from the previously published dataset ESEseq and the average effect size of member pentamers in exons from our study to (i.e. the hexamer “AGCGCA” would have member pentamers “AGCGC” and “GCGCA”) (Figure 4b). These values are well correlated (Spearman rho = 0.66) and much of the discrepancy between the two datasets are ESEseq hexamers with a value of 0, which had a non-zero value in our data.

**Figure 4).**
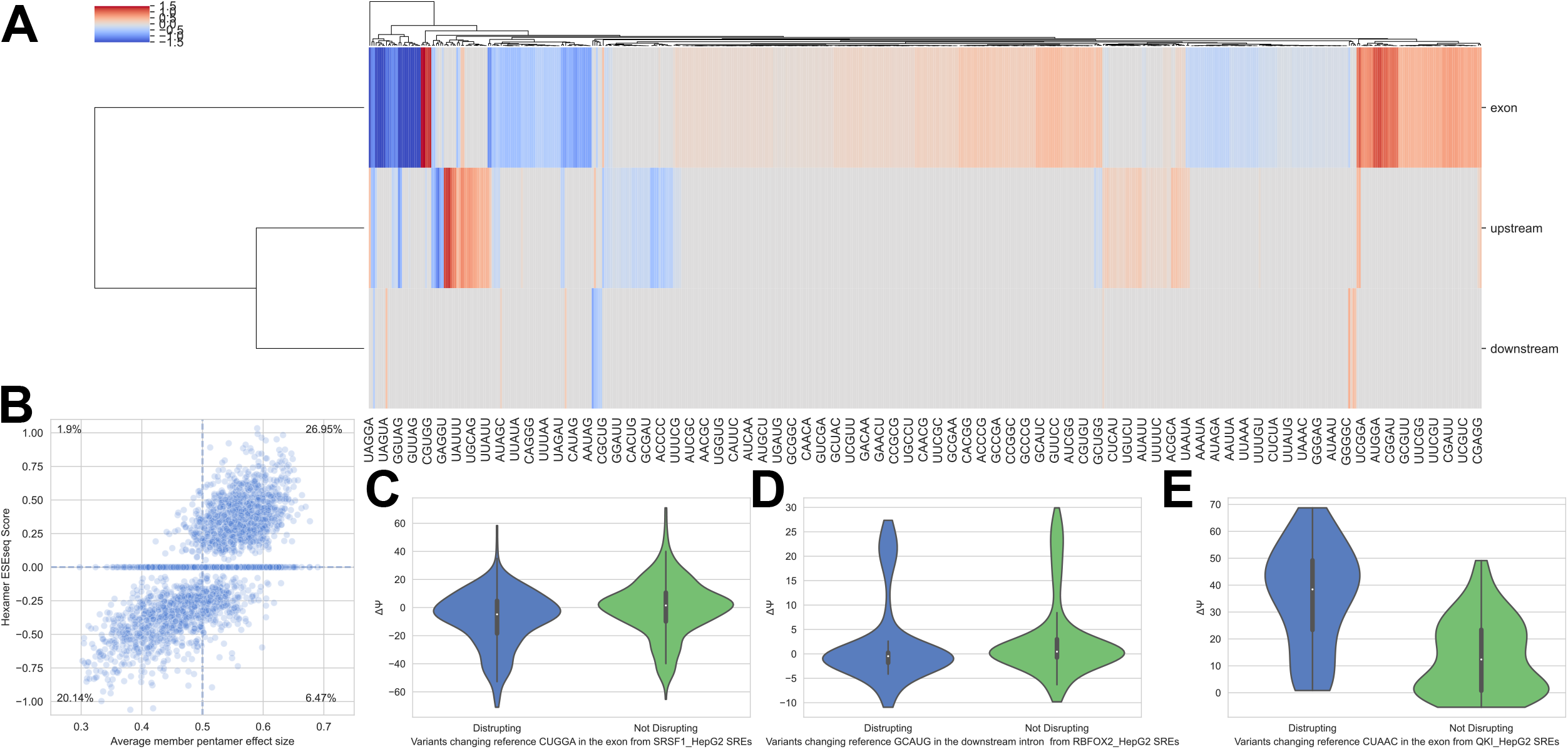
Identification of sequence elements associated with changes in splicing. **a)** A clustered heatmap of significance weighted effect size of closely related sequences containing a given pentamer, compared to those without them (-log_10_(BH corrected p-value) * (common language effect size −0.5) assessed by Mann-Whitney-U test). Data is shown for significant (BH corrected p-value ≤ 0.05) kmers only. **b)** Scatter plot showing the relationship between ESEseq hexamer score and the average member pentamer effect size from Vex-seq data. The percentages show the percentage of data points in each quadrant; remaining data lies on either the x or y axis. **c-e)** Violin plots comparing ΔΨ for sequences containing variants disrupting a kmer in the reference sequence, compared to those that do not. **c)** Violin plot for SRSF1 exonic SREs with and without CUGGA (n disrupting CUGGA = 351, n not disrupting CUGGA = 570, CLES = 0.39, BH-corrected p-value = 4.96e-07) **d)** Violin plot for RBFOX2 downstream intronic SREs with and without GCAUG (n disrupting GCAUG: 250, n not disrupting GCAUG: 427, CLES = 0.37, BH-corrected p-value = 1.04e-06) **e)** QKI exonic SREs with and without CUAAC (n disrupting CUAAC: 77, n not disrupting CUAAC: 76, CLES = 0.83, BH-corrected p-value = 6.29e-11). CLES is the common language effect size, p-values assessed by Mann-Whitney-U test and Benjamini-Hochberg multiple hypotheses correction for testing differences in loss of each 5mer in each region for each RBP dataset. All data shown from HepG2 cells.

We can extend this pentamer approach to studying the sequence requirements of SREs originating from specific RBPs in our dataset. Sequence elements associated with specific SREs would likely show a direction bias in ΔΨ if disrupted by variants. To this end, we examined whether variants which disrupted certain pentamers in specific datasets had directional bias compared to other reference sequences which were disrupted. Using this approach, we can observe pentamers associated with known motifs including CUGGA from exonic SRSF1 SREs, GCAUG from downstream intronic RBFOX2 SREs, and CUAAC in exonic QKI SREs (Figure 4 c-e). Disrupting CUGGA, a known motif for SRSF1 [27], resulted in a lower ΔΨ compared to variants which do not disrupt it, consistent with its role as and exonic splicing enhancer (Figure 4c). Using similar logic, we can make similar conclusions about GCAUG being an important sequence element for RBFOX2 as a downstream intronic enhancer, and CUAAC being an important sequence element for its role as an exonic splicing silencer mediated through QKI (Figure 4 d&e).

Understanding how variants in SREs can regulate specific exons requires a number of different pieces of information including binding, splicing behavior in absence of the RBP, and RBP motif information. To this end, we examined two splicing events in detail integrating all of these to show the influence of RBP binding on splicing regulation. First we explored a QKI binding site regulating splicing of exon 6 of the *ADGRG6* gene. This exon contained eCLIP binding sites called by both pureCLIP and iCount and showed very little read density in the input. This exon’s inclusion is increased upon *QKI* knockdown from ENCODE data, creating the hypothesis that it is acting as an ESS during normal conditions (Figure 5a top and middle). To pinpoint the location of the binding site, we used motifs from the mCross database, and scanned the exon and flanking intron sequence for motifs based on a moving average of a PSSM score. There were two local maxima of motif scores overlapping the eCLIP peaks, thus both motifs were selected for mutagenesis in Vex-seq. Sequence variants which disrupted either of these motifs increased splicing of the exon, although different variants resulted in different magnitude changes (Figure 5a bottom). While the same sequence changes in each motif result in increases in splicing, some sequence variants show different magnitude changes depending on the motif (like AAAAAC).

**Figure 5).**
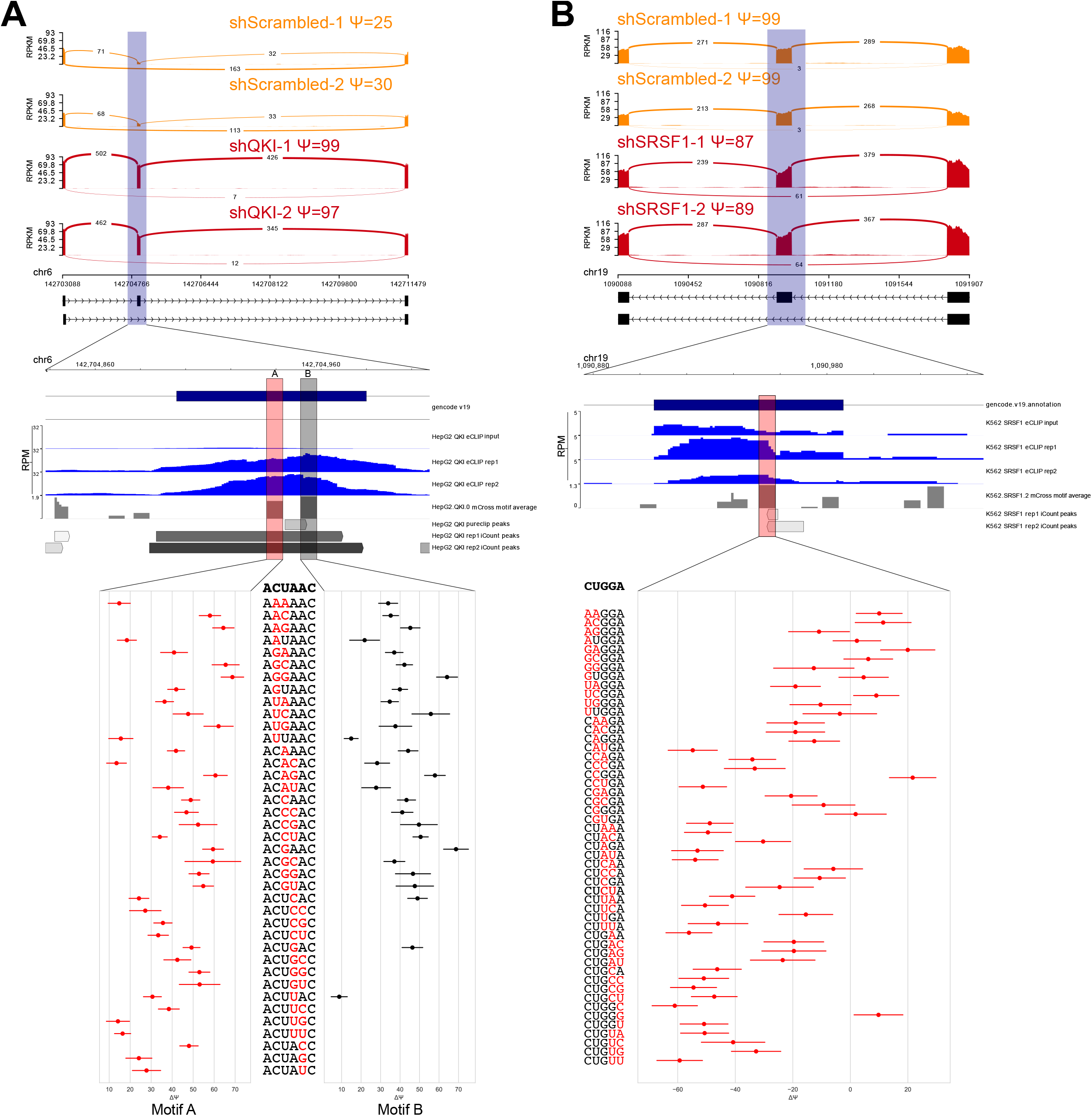
Examples identified SREs and the subsequent impacts on splicing assayed through Vex-seq. The top panel shows sashimi plots from ENCODE RBP knockdown data for scrambled and RBP knockdown. The middle panel is a zoom in on the region studied in Vex-seq which also has wiggles tracks showing ENCODE eCLIP data (input and two replicates) as well as a rolling average of mCross motifs, and peaks called by pureCLIP and iCount. The bottom panels show Vex-seq results in which the motifs for each RBP are mutated with the changed dinucleotides from the reference in red and lines represent one standard deviation. **a)** Shows a QKI SRE from HepG2 in exon 6 of *ADGRG6* with two QKI motifs which were studied by Vex-seq. **b)** Shows a SRSF1 SRE from K562 in exon 4 of *POLR2E* with one SRSF1 motif studied by Vex-seq.

Next, we decided to focus in detail on a putative SRSF1 binding site. This locus shows eCLIP peaks detected by iCount in both replicates and SRSF1 knockdown decreased exon inclusion for exon 4 of *POLR2E* in the K562 cell line (5b top and middle). These data lead to the hypothesis that SRSF1 is binding to this exon and is acting as an ESE, which is also consistent with the classical function of SRSF1 (reviewed in [35]). The highlighted region (5b middle) shows a local maximum for an mCross motif derived from K562 SRSF1 eCLIP data. Disruption of the SRSF1 binding motif in Vex-seq generally resulted in both decreases in exon inclusion or no change depending on which part of the motif was disrupted. Variants decreased ΔΨ tended to disrupt the “GGA” part of the motif, which is evident in all SRSF1 K562 derived motifs in mCross, as well as RNACompete derived SRSF1 motifs [36], and is directly bound by a domain of SRSF1 (established by nuclear magnetic resonance) [37].

While Vex-seq enables the study of many different variants and their effects on splicing, it does not maintain the context of the exon in its endogenous locus. In order to confirm that the splicing regulation we observed using Vex-seq is still relevant in the endogenous context, we used CRISPR base editors to mutate the putative motifs. For both the QKI and SRSF1 regulated exons (studied in Figure 5), we confirmed that mutating the putative binding sites did indeed change splicing. The QKI regulated *ADGRG6* exon (also shown in Figure 5a) showed that modifying either “ACUAAC” motif in the exon resulted in increased exon inclusion (Figure 6a). This is consistent with the hypothesis of the QKI binding sites acting as ESSs in the context of this exon and the Vex-seq variants disrupting these motifs also resulting in increased exon inclusion. Interestingly, the “B” base edited motif is known to be a weaker, but still used QKI motif (mCross HepG2.QKI.0 PSSM score reference motif = 10.12, variant motif score = 8.63), and shows a smaller increase in splicing relative to the “A” base edited motif, which is not predicted to be a QKI motif. It is however unknown whether the motif strength is exclusively responsible for this trend, because the same motif variants on each of the two motifs can have different magnitudes of impact on ΔΨ (Figure 5a bottom). Bulk editing worked relatively efficiently for the HepG2 samples but was less efficient for K562 samples. To enable studying these base edits in K562 cells, we used FACS to isolate K562 cells which contained the GFP-tagged base editing enzyme and grew single clones to obtain samples. These base edits targeted the putative SRSF1 binding site in the SRSF1 regulated exon 4 of *POLR2E* (similar to Figure 5b). Both edits changed the “GGA” part of the CUGGA motif studied in Figure 5b, while genotype “C” also edited nucleotides outside this sequence (Figure 6b). Each edit resulted in decreased exon inclusion, although each edit resulted in different magnitudes of splicing changes. This may reflect that while we can modify specific sequences of known motifs, there may be more subtle motif preferences outside core motifs [38], or other RBPs which may regulate variant sequences.

**Figure 6).**
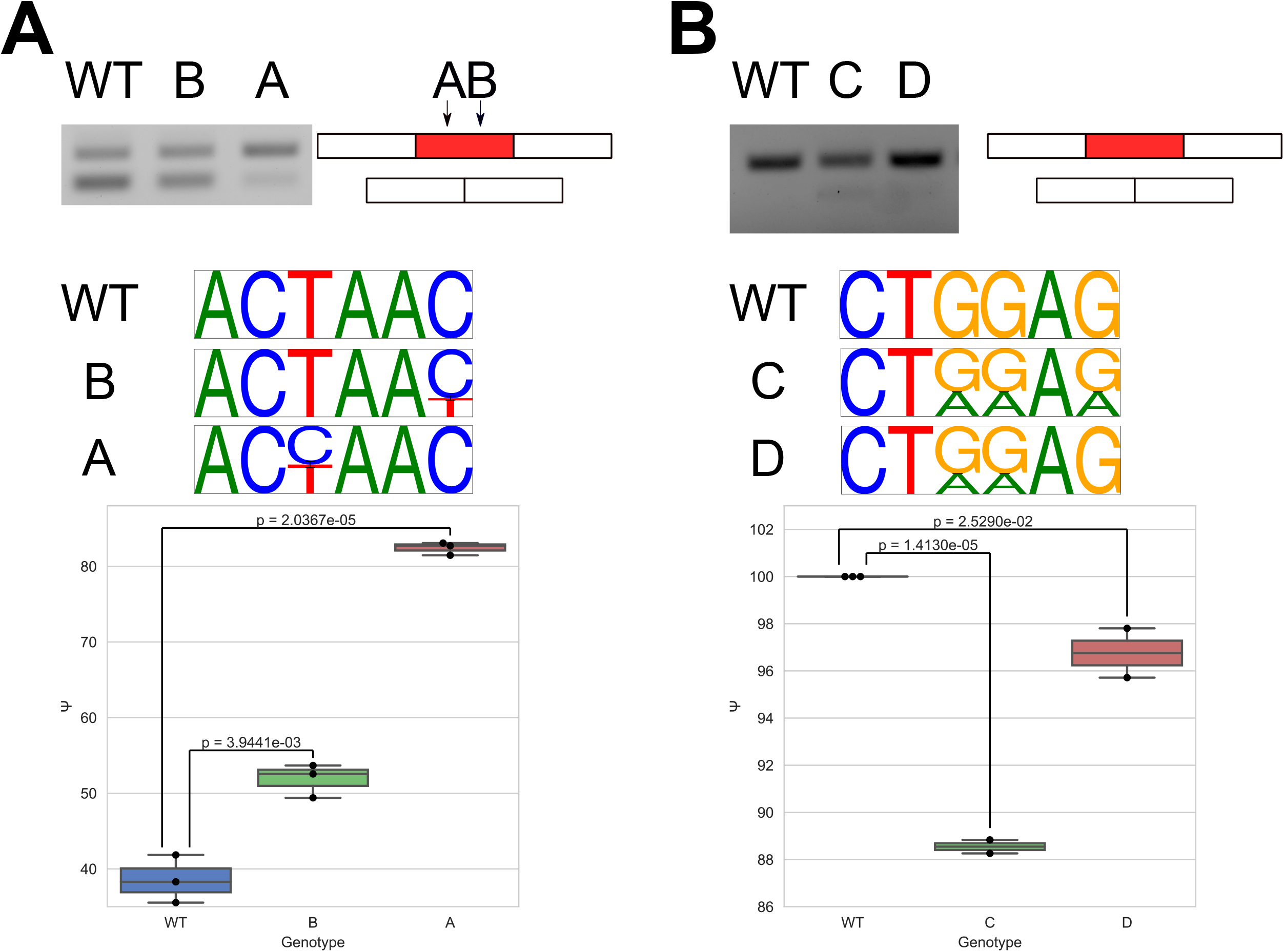
Base edits confirm effects of disrupting splicing regulatory elements. RT-PCR gels, estimated allele frequency, and splicing quantifications for **a)** the QKI regulated exon 6 of *ADGRG6* and **b)** the SRSF1 regulated exon 4 of *POLR2E.* Top shows RT-PCR gels for wildtype and base edited samples. Middle shows estimated allele frequency of sequences after base editing. Bottom shows quantification of bands in the top of the panel. Student’s t-test used to calculate p-values. A) n = 3 biological replicates each, B) n = 3 biological replicates for wildtype and 2 technical replicates for each base edited clone.

While these examples illustrate the details of splicing regulation which can be elucidated using Vex-seq, we can also observe broad trends of how variants and changes in their predicted motif strength correlate with changes in splicing. To do this, we correlated the change in motif strength the between reference and variant in the upstream intron, exon, and downstream regions and correlated it with ΔΨ. This was done by taking the log_10_ ratio of positive PSSM scores (which represent the ln(odds ratio) for a sequence deriving from the motif compared to background) between the reference and variant sequences for mCross motifs and averaging across each motif for a given RBP. For RBPamp motifs, the log_10_ ratio of RBPamp scores between the reference and variant were used as the motif scores. Significant (BH corrected p-value ≤ 0.05) Spearman correlations across each splicing event for each RBP are then binned and clustered for HepG2 and K562 (Figure 7a&b). Well characterized splicing regulation, like QKI and RBFOX2 binding to downstream introns and promoting exon inclusion, can be observed in HepG2 Figure 7a [39, 40]. Other apparent trends across cell types include SRSF1 and UCHL5 motif strength in exons being associated with increased exon inclusion (Figure 7a&b). TIA1 motif strength in downstream introns also appears to be associated with increased splicing in each cell line. Inconsistent with its known role as an ISS in upstream introns [41], PTBP1 motif strength in this region is positively correlated with exon inclusion in both cell lines. While these motif change and splicing change associations show trends observed for a number of splicing events, the most common outcome was no significant correlation and multiple hypothesis testing correction (Supplemental Figure 5a&b). Exons with significant correlations tended to have more variants and higher standard deviation of ΔΨ compared to those without significant correlations (Supplemental Figure 5c&d).

**Figure 7.**
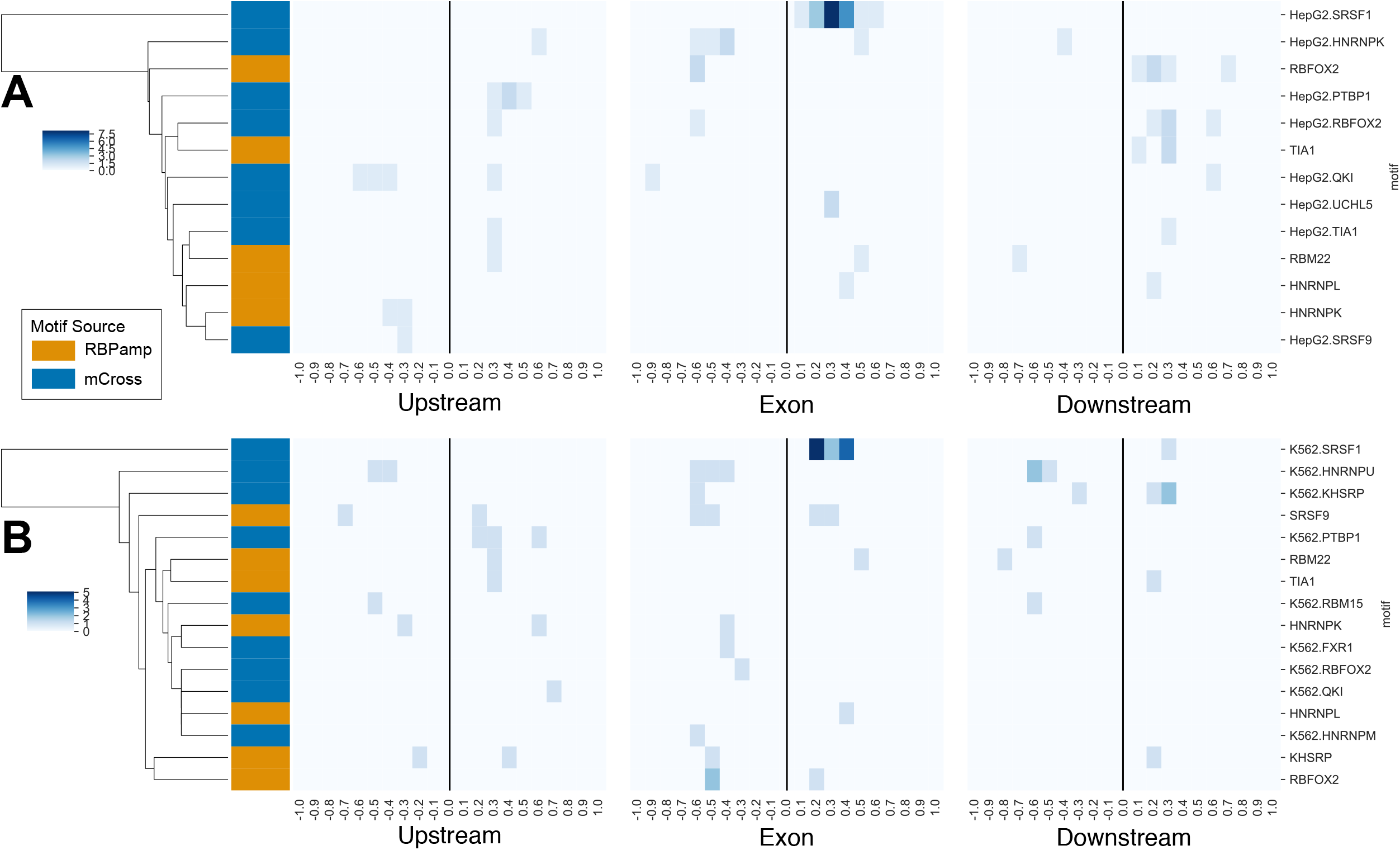
ΔΨ and motif strength changes are correlated for different RBPs at RBP-specific SREs. Clustered Spearman correlation coefficients between changes in motif strength and ΔΨ across different exons for **a)** HepG2 and **b)** K562. Only significant (BH-corrected p ≤ 0.05) correlations are shown. More intense colors reflect more exons with a correlation in that bin. Color bars show whether the motif is from mCross (blue) or RBPamp (orange).

### Validation of RBP-mediated splicing regulatory elements in external datasets

In order to ensure the splicing regulation we observed in ENCODE is not limited to K562 and HepG2 cells, we characterized the importance of splicing regulators on the splicing events in this study in other diverse cellular contexts. One hypothesis regarding the mechanism of SREs is that the extent to which they regulate splicing of a particular exon is dependent on the abundance of the RBP mediating the splicing regulation. ENCODE data has supported this hypothesis in the artificial context of RBP knockdown for the exons studied here, but it is also possible that it is true in the native context (i.e. no experimental manipulation) of cells. To investigate this, we looked for associations between RBP expression and exon inclusion in GTEx and TCGA RNA-seq datasets. Junction counts, gene expression, and metadata were extracted using Recount2 [42] from GTEx [43] and TCGA datasets. We used this data to build models predicting the Ψ of each exon that was used in this study using the expression of RBPs as predictors in order to identify RBPs with expression that is meaningfully associated with splicing of specific exons. To do this, we limited predictor genes to RBPs identified in two different RBP survey studies [44, 45] and also removed TCGA samples with missense mutations and nonsense in those genes, in order to avoid changes in splicing resulting from altered function of and RBP (i.e. SF3B1 in myelodysplastic syndromes [46]). We constructed regression models to predict logit(Ψ) of the exons in this study using the expression of RBPs as well as other available features like batch, and tissue/cancer type as predictors (see Methods for details). We also filtered out SREs which would likely yield uninformative models by removing SREs which had low diversity Ψ (top quartile – bottom quartile < 10) and filtering out SREs with fewer than 5,000 samples due to the gene harboring the exon not being expressed high enough. These resulted in 145 successful regressions. Many of the predictions performed reasonably well with a median Pearson correlation between predicted and observed Ψ on a 0 held out test set of 0.87 (0.84 median Spearman correlation) (Figure 8a). The number of gene predictors, categorical predictors, and number of samples for the models are shown in Supplemental Figure 6.

**Figure 8).**
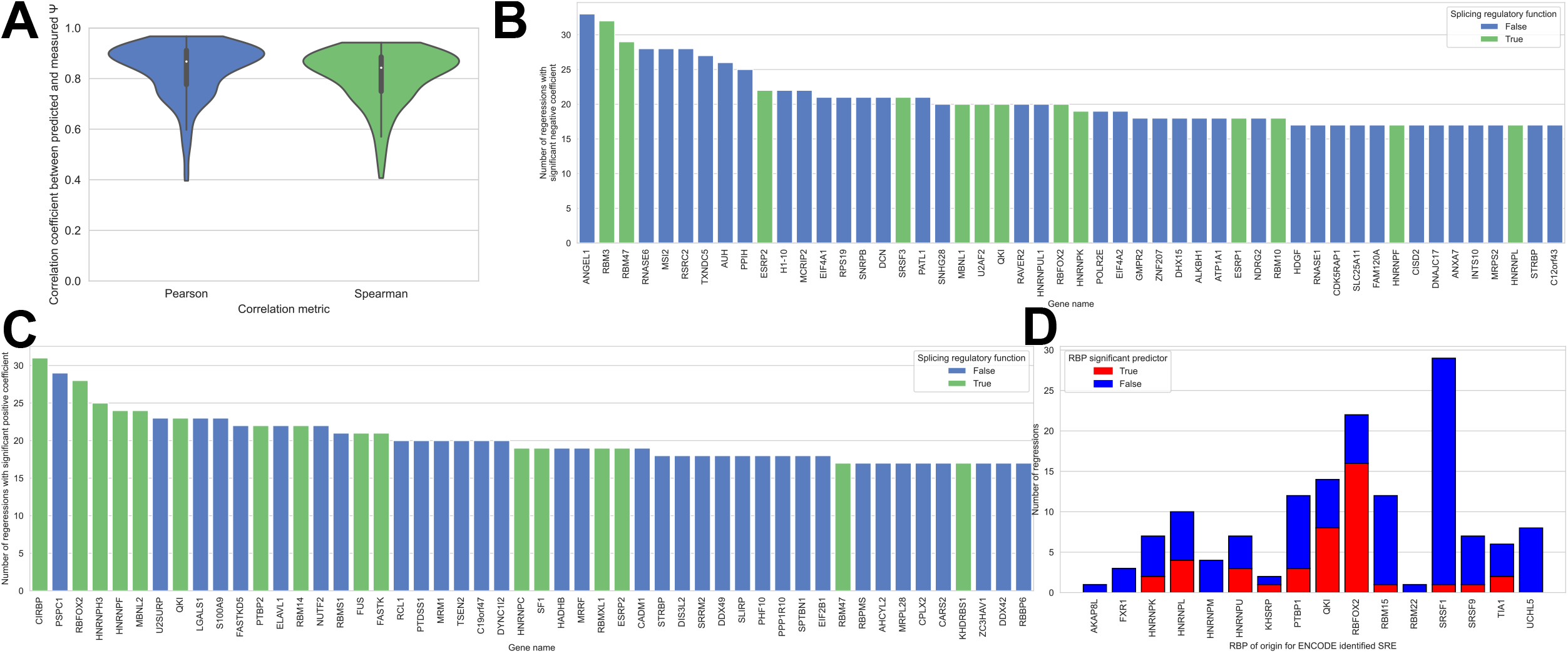
Expression of RBPs can predict Ψ of exons in this study and reveal potential splicing regulators. **a)** Pearson and Spearman correlation coefficients of predicted Ψ and real Ψ for 1/3 held out test sets for the regressions in this study. Histograms of the most frequently observed significant **b)** positive and **c)** negative gene predictors of logit(Ψ) from regression models. Known splicing regulatory proteins are shown in green. **d)** A stacked histogram showing the number of regressions for the RBPs the Vex-seq experiments were designed around. The blue shows the number of regressions where that RBP was not a significant predictor of logit(Ψ), while the red shows the number of regressions where that RBP was a significant predictor.

After constructing the models, we are able to identify the expression of which RBPs are significant predictors of splicing regulation of the exons in our study. Reassuringly, many of the top 50 most frequently observed significant predictors (which can influence splicing positively or negatively are known splicing regulators (Figure 8b&c). Focusing on the SREs in our study, we wanted to check if the RBPs with eCLIP peaks and changed splicing upon RBP knockdown were influenced by the expression of RBPs in these regression models. Indeed, many (28.97 %) of the RBPs which the Vex-seq experiments were designed to study are influenced by the expression of that RBP in other cellular contexts (Figure 8d).

## Discussion

In this study, we characterized how the splicing of exons is regulated by specific RBPs through analyzing many RBP-centric datasets from ENCODE. We identified splicing regulators in these datasets through overlapping eCLIP and KD-RNA seq in the context of HepG2 and K562 cell lines. Many RBPs showed enrichment of binding nearby exons with splicing changes compared to those without, consistent with previous studies on their roles in splicing regulation (i.e. SRSF1, RBFOX2, QKI, and HNRNPC). Other RBPs with known functions in splicing regulation however showed very little enrichment of binding near exons with splicing changes. This could be because the knockdown was not robust enough to have any consequence on splicing or other reasons not identified here. We also showed that for certain RBPs, the splicing events which changed upon RBP knockdown were more likely to have weaker binding motifs and that they were more likely to have lower co-transcriptional splicing efficiency. The sensitivity of weaker motifs to RBP knockdown reflects that when RBPs are limiting, higher affinity motifs are more likely to be bound than weaker ones, thus exons with weaker motifs are more likely to change splicing behavior. Sensitive RBPs tend to be more heavily regulated by both by redundant binding sites of a single RBP, as well as more RBPs generally binding. Sensitive events tended to have lower co-transcriptional splicing efficiency relative to robust ones. This may reflect that RBPs may need to be bound to their cognate RNA for a longer period of time in sensitive events than robust events to impart their function.

Understanding how sequence variants in splicing regulatory elements impacted splicing regulatory elements was the main focus of this paper. There were design decisions which may impact how generalizable the conclusions from this dataset are. Firstly, we only focused on SREs which were sensitive to RBP knockdown. This decision was made so that we could target high confidence SREs, however it is a limitation in that there are many more RBP binding sites near exons that were robust to knockdown in all datasets studied. These sensitive binding sites are more likely to harbor weaker motifs, multiple binding sites, or be regulated by multiple RBPs than robust ones (Figure 1b-d), which represent the majority of RBP binding sites. The other design consideration which may impact how generalizable the results are is the decision to design sequence variants around de novo motifs. When these variants were being designed, there was no motif dataset which included the majority of the RBPs of interest for this experiment (the mCross database, which contains most of these RBPs, has since been published) [27]. This may limit the specificity of the RBP-assigned motifs because the de novo motif may be a derived from another protein interacting with the RBP of interest, or non-specific co-purified RNAs which can be found in CLIP experiments [47]. Finally, the fact that these designed variants are studied in a mini-gene also limits how generalizable some of these features are, as the mini-gene removes the context of the local epigenetic environment, which has been shown to influence splicing [48, 49]. Some splicing regulatory features are impractical to study with this approach, like distal intronic splicing regulatory elements or SREs near the 5’ splice site of the upstream intron, which have been shown to have enriched binding of RBPs in alternative exons relative to constitutive ones [26]. This may be why a few reference exons didn’t predominantly splice the designed exon (Supplemental Figure 3).

Nevertheless we were able to use Vex-seq to assay the splicing behavior of sequence variants. The kmer features we identified associated with splicing regulation in each region matched up well with previously published kmers and their association with splicing in exons (Figure 4B) [50]. The presence of single kmers in reference sequences associated with known motifs for certain RBPs are significantly associated with known splicing behaviors (Figure 4 c-e), however these don’t capture the full complexity of RBP motifs, or the impact of sub-optimal motifs to influence moderate levels of splicing regulation on target exons. These subtleties are better highlighted through the examples (Figure 5), which show how variants in different SREs influence splicing regulation. Each of these highlighted interactions show how different variants in each motif have different magnitude effects on splicing. How generalizable trends in these particular SREs are to other SREs mediated by the same RBPs is unknown as these variants may have different effect sizes in different contexts, including higher RBP concentration, which can impact which RNAs are bound (Figure 1b) [51].

When observing correlations between changes in motif strength and changes in splicing, we observe a number of both expected and unexpected behaviors when compared prior literature and analysis of KD-RNA seq eCLIP overlaps. QKI appears to be an ISE when binding in the downstream intron but suppress exon inclusion when binding in the upstream intron, consistent with prior literature [40, 52] (Figures 7a, Figure 1a). Less evidence is available concerning its role as an ESS, which is characterized in our analysis (Figures 5a and 6a). We also have identified a UCHL5 as a potential novel splicing regulator (Figures 1a, 7a), which has also recently been implicated in splicing regulation based on other analyses of these same datasets [53, 54]. Direct interactions between RNA and UCHL5, a deubiquitylase without known RNA-binding domains, may be an unexpected mechanism of splicing regulation, however Carazo et al., notes that UCHL5 may have protein interactions with SRSF1 [54]. UCHL5 has also been shown to interact with SNRNPA (a component of U1 snRNP) through high-throughput fractionation mass spectrometry experiments, reflecting that it is possible it acts to regulate splicing through post-translationally modifying other known splicing regulators [55]. The observation in our data that exons without meaningful correlations between motif strength and splicing changes tend to have fewer designed variants (Supplemental Figure 5c) highlights that there is likely more detailed splicing regulation that can be studied in these exons with more comprehensive mutagenesis.

Finally, we analyzed other datasets external to ENCODE and observed that many of the splicing regulatory elements that we identified have relationships between RBP expression and splicing in those datasets. We observed that 28.97 % of SREs highlighted in this study showed the RBP of interest as a significant predictor of splicing. This analysis also shows how other RBPs which have not yet been assayed in ENCODE may be important splicing regulators. Leveraging large public RNA-seq datasets to gain information about splicing changes is not conceptually new [56], however, to our knowledge its application to directly identify splicing regulators is. There are of course many caveats with this analysis approach, however the fact that many of the most frequently observed significant predictors are splicing factors, the reasonable correlation between predictions and a held-out test set, and that many ENCODE SREs validate is reassuring that these models may yield some insights. Some biological caveats include that not all splicing events have diverse enough Ψ to yield meaningful results, and that RNA expression (especially aggregated at the gene level) of an RBP is not always a reliable indicator of protein abundance or functionality. Other biological caveats are that RBP paralogs may compensate for each other at certain binding sites and that the expression of certain genes are likely correlated. Additionally, there are technical confounders that include that not all samples could reliably be assigned to a sequencing batch. Furthermore, a histone mRNA that is not typically polyadenylated is a frequently observed significant predictors of Ψ, suggesting that variable efficiency in mRNA purification may be an unseen technical confounder which is impacting the analysis. Nevertheless, RBPs frequently shown to be meaningful predictors of splicing, which are not currently characterized may be avenues of interest for researchers investigating splicing regulation. While we only investigated splicing events that were altered with protein knockdown, this regression approach may be useful for other splicing events in the future.

This study has shed light on the role of RBP binding sites in splicing regulation, however further experiments in the context of the genome should be done in order to assay RBP binding sites and the effect of altering them more accurately. These experiments however are labor intensive at the scale that this paper studies using Vex-seq. While the sequence variants studied here are designed to disrupt SREs, uncovering the impact of genetic variants found in human populations on splicing regulation are likely to be fruitful avenues of research.

## Conclusions

In this study, we have identified splicing regulators in ENCODE datasets, and have characterized features important for splicing regulation in the context of RBP depletion. Exons which tend to be sensitive to RBP depletion tend to be mediated by weaker motifs and more binding sites. Furthermore, these exons also tend to bound my more RBPs and spliced less efficiently at steady state, relative to exons which do not change splicing upon RBP depletion, but are still bound by that RBP. We then go on to identify RBP-agnostic and RBP-specific sequence features associated with splicing regulation using Vex-seq, and show that across some exons, that *in silico* motif strength correlates with splicing strength at RBP-regulated exons. We also validate that perturbation of a small subset of these splicing regulatory elements in the context of the genome changes splicing in a manner consistent with Vex-seq mutations. We additionally validate some of these SREs by association in the context of the GTEx and TCGA samples and identify other potential splicing regulators not identified in ENCODE data. These results shed light on some of the principles underlying splicing regulation by RBPs mediated by SREs and identify potentially uncharacterized splicing regulators.

## Methods

### eCLIP and KD RNA-seq data processing

Splicing regulatory and novel RBPs were identified based on the annotations in [26]. eCLIP bam files aligned to hg19 were downloaded from www.encodeproject.org and processed to identify binding sites through iCount and pureCLIP pipelines [57]. For pureCLIP peak calling, read headers from downloaded bam files were reformatted such that they could be deduplicated with UMI-tools (v1.0.0) [58]. After deduplication, common crosslink motifs were downloaded from (https://raw.githubusercontent.com/skrakau/PureCLIP_data/master/common_CL-motifs/dreme.w10.k4.xml) and instances were identified with compute_CLmotif_scores.sh (provided as part of pureCLIP). The pureCLIP (v1.3.1) HMM was trained on chromosomes 1, 2, and 3 and peaks were called jointly on the bam files of each biological replicate incorporating common crosslink motifs and size matched input.

For iCount (v2.0.0) peak calling (T. Curk, et al., manuscript in preparation, available at https://github.com/tomazc/iCount) UMIs in read names were reformatted such that they would be compatible with iCount cDNA quantification. iCount xlsites was used with the parameters -- group_by star and --quant cDNA, then iCount peaks and clusters were used to identify clusters of binding sites. Peaks on chrM for both pipelines were removed.

rMATs pre-processed data for KD RNA-seq was downloaded from www.encodeproject.org.

### Splicing regulatory element identification and characterization

To identify splicing regulatory elements, we used bedtools (v2.27.1) and pybedtools (v0.8.0) to intersect peaks identified with iCount with each splicing event quantified from the rMATS files [59, 60]. Peaks used for this analysis were called on each replicate separately, then merged using bedtools merge. For the upstream and downstream introns, we considered 300 nts up or downstream from the exon boundaries. Upregulated and downregulated events identified by having FDR-corrected p-value ≤ 0.05 and |Inclusion Level Difference| ≥ 0.01, all events not meeting these criteria were considered background events. Fisher’s exact test was used to identify binding events which were enriched in either upregulated or downregulated splicing events within the upstream intron, exon, or downstream intron. Multiple hypothesis testing correction for comparisons for each region and splicing direction was done using Benjamini-Hochberg correction.

Sequences assessed for peak strength were expanded by 30 nts on each side using and strand specific sequences were retrieved using bedtools slop and getfasta [59]. For this analysis, we considered “sensitive” as any splicing event that changes with RBP knockdown (FDR-corrected p-value ≤ 0.05 and |Inclusion Level Difference| ≥ 0.01) with a peak within 300 nts of the exon and “robust” as the same binding patterns around exons which do not change with RBP knockdown. The scoring of mCross motifs was done though retrieving TRANSFAC files from mCross (http://zhanglab.c2b2.columbia.edu/data/mCross/eCLIP_mCross_PWM.tgz) [27]. Position specific scoring matrices were calculated under a uniform nucleotide distribution and scores were calculated for each of the sequences using the “motifs” module in Biopython [61]. The sum of the positive scores was calculated for each motif identified for a dataset and were averaged. If an mCross motif was not available in a specific cell line, the motif for the same RBP in the other cell line was used. RBPamp (Jens, M. et al., submitted) motifs were calculated based on RNA bind-n-seq data [24, 26]. Average scores across each sequence were calculated using the eval_model function within RBPamp (interpreted as 1/Kd for a given sequence).

For the co-transcriptional splicing efficiency analysis, co-transcriptional splicing efficiency was downloaded from Table S1 from [32]. Intron coordinates from this study were matched to intron coordinates from rMATs events. If available, the mean co-transcriptional splicing efficiency of each flanking intron was used as the co-transcriptional splicing efficiency of the event, if only one was available, that was used instead of the mean. If both introns were missing from the table, the events were ignored as not all introns from rMATs were in the table due to the presence of novel splicing events in the rMATs datasets and robustness thresholds for identifying co-transcriptional splicing efficiencies set by the authors which provided the data. For all sensitive vs. robust comparisons, datasets were excluded from the analysis if they did not contain more than 10 sensitive events.

### Vex-seq design

eCLIP peaks from each peak caller were used as inputs to de novo motif identifiers DREME (v5.0.5 with parameters -maxk 6 -norc) and ssHMM (v1.0.5 with default parameters) [62, 63]. Peaks from each peak caller for sensitive events for exons < 100nts were manually associated with the nearest local maxima of motifs to identify candidate regions for mutagenesis. Saturating di-nucleotide mutagenesis was designed over motif local maxima nearest to the peak. At least 50 nts of the upstream intron and at least 20 nts of the downstream intron were included in the design. For peaks in exons, leftover sequences (until the tested sequence made up 169-170 nts) were evenly distributed to upstream and downstream introns. For peaks in either intron, excess leftover sequence was assigned to that intron. Common sequences required for proper assembly splicing reporter as well as the barcodes were added to test sequences. Sequences were designed as with a common 5’ primer, the test sequence, common MfeI and SpeI sites, barcode, and common 3’ primer with XbaI site. Common sequences are as follows: common 5’ primer CTGACTCTCTCTGCCTC, common MfeI and SpeI sites CAATTGACTACTAGT, and common 3’ primer TCTAGAGGGCCCGTTTA. 10mer barcodes were generated using the R package DNABarcodes, such that each barcode had a Hamming distance of 2 from other barcodes and did not contain any restriction enzyme sites used in the plasmid assembly [64]. Three barcodes are associated with each test sequence. These 230mers were then synthesized using array-based DNA synthesis from Agilent. These sequences are available in Additional file 2.

### Vex-seq library assembly and sequencing

The Vex-seq splicing reporter library was assembled as previously described with some minor modifications [15]. To reduce the number of transcripts splicing to the plasmid backbone, we changed the way that the library was first cloned in. Modified pcAT7-Glo1 was linearized by digestion with PstI and XbaI, the ends were then blunted using DNA Polymerase I, (Large Klenow) Fragment (to remove the AG in the PstI overhang which could be used as a 3’ splice site). The synthesized oligonucleotides were then inserted into the linearized plasmid using Gibson assembly [65]. The rest of the plasmid library construction was the same as previously described.

Sequencing library construction was performed as previously described [15] with one minor change. In order to increase the diversity of UMIs in our library, we changed the reverse transcription primer to GTGACTGGAGTTCAGACGTGTGCTCTTCCGATCTNNNNNNNNNNGCTGATCAGCGGGTTT AAACG. The resulting library was sequenced on a NextSeq 550 using a 300 cycle high-output kit with a 75 nt read 1 and a 225 nt read 2. Modified pcAT7-Glo1 is available through Addgene (Plasmid #160996). A detailed protocol, as well as a table with primer sequences used in Vex-seq library construction and sequencing is provided in supplemental methods.

### Cell culture and transfections

For Vex-seq transfections, HepG2 cells were grown in 6-well plates to 0.7×10^6^ cells/well. Cells were then transfected with 5μg of plasmid DNA (FuGENE HD Transfection Reagent). K562 cells were grown to a density of 1×10^6^ cells and electroporated with 5μg of plasmid DNA *(Amaxa^®^* Cell Line Nucleofector^®^ Kit V, Lonza). Cells from each cell line were cultured for an additional 48 hours until total RNA isolation (Maxwell RSC simplyRNA kit, Promega).

For CRISPR base editing experiments, HepG2 cells were grown in 6-well plates to 0.7×10^6^ cells/well. HepG2 cells were then transfected with 1,250ng gRNA plasmid and 3,750ng of either pCMV_AncBE4max_P2A_GFP or pCMV_ABEmax_P2A_GFP [66] (FuGENE HD Transfection Reagent Cat#E2311, Promega). HepG2 cells were cultured for an additional 48 hours then green cells were isolated using flow cytometry (Aria II). K562 cells were grown to 1×10^6^ and transfected with 1,000ng of gRNA plasmid and 3,000 ng of pCMV_AncBE4max_P2A_GFP (*Amaxa*^®^ Cell Line Nucleofector^®^ Kit V, Lonza). K562 cells were cultured for an additional 48 hours then single clones of green cells were isolated using flow cytometry (Aria II). These single clones were cultured for 4 weeks prior to harvesting of genomic DNA and RNA. Genomic DNA from each cell line was isolated (QuickExtract DNA Extraction Solution, Lucigen). Total RNA was isolated from each cell line (Maxwell RSC simplyRNA kit, Promega). pCMV_ABEmax_P2A_GFP was a gift from David Liu (Addgene plasmid # 112101; http://n2t.net/addgene:112101; RRID:Addgene_112101). pCMV_AncBE4max_P2A_GFP was also a gift from David Liu (Addgene plasmid # 112100; http://n2t.net/addgene:112100; RRID:Addgene_112100).

### Vex-seq data processing

After demultiplexing UMIs were extracted from read 2 using UMItools (v1.0.0) extract (with --bc-pattern=NNNNNNNNNN) [58]. Barcodes were extracted using custom python scripts to identify the reference which the sequences the sequences should be aligned to. Reads were aligned to their barcode associated reference with hisat2 [67] (v2.1.0) with parameters -no-softclip --dta-cufflinks. Read pairs were then deduplicated based on the mapping coordinates and the UMI using UMItools (v1.0.0) dedup. To identify novel splice sites, we used stringtie (v2.0.3) [33] on all the alignments for a particular sequence and merged them. Properly paired reads were extracted from the bam files and splice junctions for each read pair were extracted using pysam (v0.15.4) (https://github.com/pysam-developers/pysam) [68] and counted for all junctions identified with stringite. These junction counts were then scaled to TPM before using SUPPA2 (v2.3) [34] psiPerEvent and diffSplice (with --method empirical) to identify differential splicing.

For plasmid sequencing data processing, reads were aligned to a pooled plasmid reference using BWA-MEM (v0.7.17). For the primary plasmid pool sequencing samples, the BWA parameter “-I 240,24,265,225” was applied and for the final plasmid pool “-I 298,29,315,275” was applied. Concordantly mapped read pairs were retained for variant calling. Variants were identified using samtools mpileup and bcftools [69] (samtools and htslib v1.9) and variant frequency was identified by dividing AD by DP in the resulting vcf file.

### Base editing and RT-PCR

All gRNAs for base edits were designed using BE-Designer [70] and cloned into MLM3636. Briefly, the 5’ and 3’ oligos encoding the guide sequence, were hybridized, and subcloned into BsmbI digested MLM3636. Plasmids were sanger sequenced to ensure the correct insert was contained. gRNA plasmids were used and genomic DNA and total RNA was isolated as described in the “Cell culture and transfections” section. Genomic DNA was PCR amplified and sanger sequenced to identify the success and efficiency of edits. Base editing efficiency was estimated using Tracy [71]. RNA isolation, cDNA was synthesized using SuperScript III and random hexamer primers. RT-PCR amplicons were electrophoresed on 2 % agarose gels and imaged on a ChemiDoc MP (Bio-Rad). The resulting images were analyzed using ImageJ (v1.52a), by subtracting background in a 30 pixel radius and calculating the intensity of bands associated with the included or excluded isoform. Inclusion was then calculated as the (intensity of the included band x 100) / (the intensity of the included band + intensity of excluded band). All primers, and their applications relevant to the base editing experiments are listed in Additional file 6. MLM3636 was a gift from Keith Joung (Addgene plasmid # 43860; http://n2t.net/addgene:43860; RRID:Addgene_43860).

### Motif analysis

For all splicing events (previously annotated and unannotated), we scored motifs from RBPamp and mCross motifs as described in the “Splicing regulatory element identification and characterization” section across the upstream intron, exon and downstream intron. We then correlated ΔΨ to the log_10_(variant motif score in a region/ reference motif score in a region) for all variants designed around an SRE for a specific RBP in that cell line. The spearman correlation coefficients across each region were then clustered for each cell line. Correlation p-values in each region were adjusted for multiple hypothesis testing using Benjamini-Hochberg correction. Correlation coefficients which were not significant (BH corrected p-value ≤ 0.05) were plotted as 0 (Figure 7). For examples highlighted in Figure 5, pygenometracks [72] was used to create the parts of the figure depicting genomic data (i.e. gene models, eCLIP data, motif data), and rmats2sashimiplot (https://github.com/Xinglab/rmats2sashimiplot) was used to create sashimi plots.

### GTEx TCGA splicing regressions

RBPs considered as predictors were identified from multiple RBP census studies [44, 45]. In order to identify TCGA samples which contained missense or nonsense mutations in the considered RBPs, we used TCGAbiolinks [73] to identify tumor samples with these mutations as identified by the “mutect” pipeline using the GDCquery_Maf function. Gene expression and junction counts files for TCGA and GTEx (SRP012682) were downloaded from Recount2 [42]. Splice junctions corresponding to SREs were identified using custom python scripts. Categorical variables corresponding for samples were parsed from Recount2 retrieved metadata including batch (“smgebtch” for GTEx and “cgc_case_batch_number” for TCGA), tissue/tumor type, and project (GTEx or TCGA). Samples with fewer than 10 total reads supporting either included or excluded isoforms were removed from analysis. Exon inclusion rate was calculated by dividing the included junction counts by included junction counts plus excluded junction counts x 2. Gene expression values were calculated by dividing the gene counts by the gene length and dividing by the length normalized gene counts for all genes. These gene expression values were then transformed into Z-scores for each gene across all samples and this was used as input for regressions. To identify meaningful variables, we used glmnet (v3.0.2) [74] in R (v3.6.0) to train a model to predict logit(Ψ) using 10-fold cross validation in an elastic net regression using junction counts for the event in interest as sample weight (using the cv.glmnet function with alpha = 0.5). Non-zero gene coefficients under the best lambda parameter were extracted and used as inputs (as well as categorical predictors) to a conventional regression model using the glm function with a 1/3 held out test set. Significance of predictors was adjusted for by using Benjamini-Hochberg multiple hypothesis testing correction. Splicing regulators as discussed in this section were identified by combining splicing regulators identified in [26] with splicing regulators identified under the gene ontology identifier GO:0043484 using AmiGO [75–77].

## Declarations

### Ethics approval and consent to participate

Not applicable

### Consent for publication

Not applicable

### Availability of data and materials

Fastq files as well as summary files for the data generated in this study are available at GEO under the accession GSE164752. The modified pcAT7-Glo1 plasmid is available from Addgene (#160996). Code producing figures, as well as example scripts for data pre-processing can be found on github (https://github.com/scottiadamson/Splicing_regulatory_elements_paper). Additional processed data used as inputs for figures that are large and not available on github or as additional files are available on Zenodo (https://doi.org/10.5281/zenodo.4681667).

### Authors’ contributions

BRG and SIA conceived of the experiments. SIA designed oligos, built Vex-seq libraries, and performed all data analysis in this manuscript. LZ performed cell culture, transfection, and flow cytometry experiments. SIA and BRG wrote the manuscript. All authors read and approved the final manuscript.

## Acknowledgements

We would like to thank the RNA binding protein group of the ENCODE consortium for generating and making publicly accessible all of the data used in this paper. Additionally, we would like to thank Marvin Jens and Chris Burge for giving us early access to RBPamp ahead of publication. We would also like to thank members of the Graveley lab and members of the RNA binding protein group of the ENCODE consortium for feedback and constructive criticism on this work. We would also like to thank the authors of Recount2 for processing GTEx and TCGA data and making it publicly available, as well as all of the patient donors, scientists and funders that made these datasets possible and available.

## Supplemental Figure Legends

<u>**Supplemental Figure 1.** Distribution of testable and all identified SREs from the ENCODE datasets</u>. Histograms showing the distribution exons bound by specific RBPs which change splicing upon RBP knockdown (sensitive events). Histogram shows RBPs which **a)** decrease inclusion upon RBP knockdown and **b)** RBPs which increase inclusion upon RBP knockdown. Testable exons (≤101 nts with an RBP peak overlapping a testable region) are shown in red, and the remainder are shown in blue.

<u>**Supplemental Figure 2.** Quality control for the Vex-seq sequence design and splicing behavior.</u> **a)** A histogram of the representation of each of the barcodes in the final plasmid pool. **b)** A histogram of the variant frequency observed for each barcode. The dotted line represents the threshold of variant frequency for which barcodes were excluded from future analysis. For **c** and **d**, above the diagonal shows the Pearson and Spearman correlation coefficients, the diagonal has histograms describing the distribution of values for each sample, and below the diagonal are scatter plots. **c)** Correlations of Ψ values between biological samples. **d)** Correlations of ΔΨ values between biological samples.

<u>**Supplemental Figure 3.** Ψ of annotated and unannotated exons with reference sequences in Vex-seq.</u> Boxplots showing the Ψ values of reference sequence exons with unannotated exons detected as well as the annotated exons for the same sequences for **a)** HepG2 and **b)** K562.

<u>**Supplemental Figure 4.** Number and categories of A5SS events identified in Vex-seq.</u> Violin plots showing the distribution of ΔΨs for significantly changing alternative 5’ splice site splicing events evaluated by SUPPA2 broken down by **a)** short and **b)** long 5’ splice site changes as evaluated by MaxEntScan. Histograms above the plots show the number of events in each category.

<u>**Supplemental Figure 5.** Properties of SREs with and without meaningful motif strength change and ΔΨ correlations.</u> Stacked bar charts showing the number of correlations between motif strength and ΔΨ across exons in **a)** HepG2 and **b)** K562 cell lines. Violin plots showing the **c)** number of observations (p = 7.7e-08, CLES = 0.66) and **d)** standard deviation of ΔΨ (p = 2.02e-4, CLES = 0.61) between exons with significant and not significant correlations between change motif strength with ΔΨ. For C and D, n significant = 122, n not significant = 415, statistics assessed by Mann-Whitney-U; CLES is the common language effect size.

<u>**Supplemental Figure 6.** Number of samples and predictors used as inputs to regression models</u>. Histograms showing the distribution of **a)** the number of samples, **b)** the number of categorical predictors, and **c)** the number of gene predictors for the regressions predicting Ψ of exons regulated SREs in this study using GTEx and TCGA data.

## Citations

1. Cingolani P, Platts A, Coon M, Nguyen T, Wang L, Land SJ, et al. A program for annotating and predicting the effects of single nucleotide polymorphisms, SnpEff: SNPs in the genome of Drosophila melanogaster strain w1118; iso-2; iso-3. Fly. 2012;6:80–92.

2. Hinrichs AS, Raney BJ, Speir ML, Rhead B, Casper J, Karolchik D, et al. UCSC Data Integrator and Variant Annotation Integrator. Bioinformatics. 2016;32:1430–2.

3. McLaren W, Gil L, Hunt SE, Riat HS, Ritchie GRS, Thormann A, et al. The Ensembl Variant Effect Predictor. Genome Biol. 2016;17:122.

4. Wang K, Li M, Hakonarson H. ANNOVAR: functional annotation of genetic variants from high-throughput sequencing data. Nucleic Acids Research. 2010;38:e164–e164.

5. Manning KS, Cooper TA. The roles of RNA processing in translating genotype to phenotype. Nat Rev Mol Cell Biol. 2017;18:102–14.

6. Scotti MM, Swanson MS. RNA mis-splicing in disease. Nat Rev Genet. 2016;17:19–32.

7. Li YI, van de Geijn B, Raj A, Knowles DA, Petti AA, Golan D, et al. RNA splicing is a primary link between genetic variation and disease. Science. 2016;352:600–4.

8. Singh RN, Singh NN. Mechanism of Splicing Regulation of Spinal Muscular Atrophy Genes. Adv Neurobiol. 2018;20:31–61.

9. Xiong HY, Alipanahi B, Lee LJ, Bretschneider H, Merico D, Yuen RKC, et al. RNA splicing. The human splicing code reveals new insights into the genetic determinants of disease. Science. 2015;347:1254806.

10. Xiong HY, Barash Y, Frey BJ. Bayesian prediction of tissue-regulated splicing using RNA sequence and cellular context. Bioinformatics. 2011;27:2554–62.

11. Leung MKK, Xiong HY, Lee LJ, Frey BJ. Deep learning of the tissue-regulated splicing code. Bioinformatics. 2014;30:i121-129.

12. Cheng J, Nguyen TYD, Cygan KJ, Celik MH, Fairbrother WG, Avsec Z, et al. MMSplice: modular modeling improves the predictions of genetic variant effects on splicing. Genome Biol. 2019;20:48.

13. Jaganathan K, Kyriazopoulou Panagiotopoulou S, McRae JF, Darbandi SF, Knowles D, Li YI, et al. Predicting Splicing from Primary Sequence with Deep Learning. Cell. 2019;176:535–548.e24.

14. Rentzsch P, Schubach M, Shendure J, Kircher M. CADD-Splice—improving genome-wide variant effect prediction using deep learning-derived splice scores. Genome Medicine. 2021;13:31.

15. Adamson SI, Zhan L, Graveley BR. Vex-seq: high-throughput identification of the impact of genetic variation on pre-mRNA splicing efficiency. Genome Biol. 2018;19:71.

16. Soemedi R, Cygan KJ, Rhine CL, Wang J, Bulacan C, Yang J, et al. Pathogenic variants that alter protein code often disrupt splicing. Nat Genet. 2017;49:848–55.

17. Cheung R, Insigne KD, Yao D, Burghard CP, Wang J, Hsiao Y-HE, et al. A Multiplexed Assay for Exon Recognition Reveals that an Unappreciated Fraction of Rare Genetic Variants Cause Large-Effect Splicing Disruptions. Mol Cell. 2019;73:183–194.e8.

18. Rosenberg AB, Patwardhan RP, Shendure J, Seelig G. Learning the sequence determinants of alternative splicing from millions of random sequences. Cell. 2015;163:698–711.

19. Mikl M, Hamburg A, Pilpel Y, Segal E. Dissecting splicing decisions and cell-to-cell variability with designed sequence libraries. Nat Commun. 2019;10:4572.

20. Baeza-Centurion P, Miñana B, Schmiedel JM, Valcárcel J, Lehner B. Combinatorial Genetics Reveals a Scaling Law for the Effects of Mutations on Splicing. Cell. 2019;176:549–563.e23.

21. Ke S, Anquetil V, Zamalloa JR, Maity A, Yang A, Arias MA, et al. Saturation mutagenesis reveals manifold determinants of exon definition. Genome Res. 2018;28:11–24.

22. Wong MS, Kinney JB, Krainer AR. Quantitative Activity Profile and Context Dependence of All Human 5’ Splice Sites. Mol Cell. 2018;71:1012–1026.e3.

23. Julien P, Miñana B, Baeza-Centurion P, Valcárcel J, Lehner B. The complete local genotype-phenotype landscape for the alternative splicing of a human exon. Nat Commun. 2016;7:11558.

24. Lambert N, Robertson A, Jangi M, McGeary S, Sharp PA, Burge CB. RNA Bind-n-Seq: quantitative assessment of the sequence and structural binding specificity of RNA binding proteins. Mol Cell. 2014;54:887–900.

25. Van Nostrand EL, Pratt GA, Shishkin AA, Gelboin-Burkhart C, Fang MY, Sundararaman B, et al. Robust transcriptome-wide discovery of RNA-binding protein binding sites with enhanced CLIP (eCLIP). Nat Methods. 2016;13:508–14.

26. Van Nostrand EL, Freese P, Pratt GA, Wang X, Wei X, Xiao R, et al. A large-scale binding and functional map of human RNA-binding proteins. Nature. 2020;583:711–9.

27. Feng H, Bao S, Rahman MA, Weyn-Vanhentenryck SM, Khan A, Wong J, et al. Modeling RNA-Binding Protein Specificity In Vivo by Precisely Registering Protein-RNA Crosslink Sites. Mol Cell. 2019;74:1189–1204.e6.

28. Drexler HL, Choquet K, Churchman LS. Splicing Kinetics and Coordination Revealed by Direct Nascent RNA Sequencing through Nanopores. Mol Cell. 2020;77:985–998.e8.

29. Pandya-Jones A, Black DL. Co-transcriptional splicing of constitutive and alternative exons. RNA. 2009;15:1896–908.

30. Fong N, Kim H, Zhou Y, Ji X, Qiu J, Saldi T, et al. Pre-mRNA splicing is facilitated by an optimal RNA polymerase II elongation rate. Genes Dev. 2014;28:2663–76.

31. Dujardin G, Lafaille C, de la Mata M, Marasco LE, Muñoz MJ, Le Jossic-Corcos C, et al. How slow RNA polymerase II elongation favors alternative exon skipping. Mol Cell. 2014;54:683–90.

32. Saldi T, Riemondy K, Erickson B, Bentley DL. Alternative RNA structures formed during transcription depend on elongation rate and modify RNA processing. Mol Cell. 2021.

33. Pertea M, Pertea GM, Antonescu CM, Chang T-C, Mendell JT, Salzberg SL. StringTie enables improved reconstruction of a transcriptome from RNA-seq reads. Nature Biotechnology. 2015;33:290–5.

34. Trincado JL, Entizne JC, Hysenaj G, Singh B, Skalic M, Elliott DJ, et al. SUPPA2: fast, accurate, and uncertainty-aware differential splicing analysis across multiple conditions. Genome Biol. 2018;19:40.

35. Das S, Krainer AR. Emerging functions of SRSF1, splicing factor and oncoprotein, in RNA metabolism and cancer. Mol Cancer Res. 2014;12:1195–204.

36. Ray D, Kazan H, Cook KB, Weirauch MT, Najafabadi HS, Li X, et al. A compendium of RNA-binding motifs for decoding gene regulation. Nature. 2013;499:172–7.

37. Cléry A, Sinha R, Anczuków O, Corrionero A, Moursy A, Daubner GM, et al. Isolated pseudo-RNA-recognition motifs of SR proteins can regulate splicing using a noncanonical mode of RNA recognition. Proc Natl Acad Sci U S A. 2013;110:E2802–11.

38. Cléry A, Krepl M, Nguyen CKX, Moursy A, Jorjani H, Katsantoni M, et al. Structure of SRSF1 RRM1 bound to RNA reveals an unexpected bimodal mode of interaction and explains its involvement in SMN1 exon7 splicing. Nature Communications. 2021;12:428.

39. Yeo GW, Coufal NG, Liang TY, Peng GE, Fu X-D, Gage FH. An RNA code for the FOX2 splicing regulator revealed by mapping RNA-protein interactions in stem cells. Nat Struct Mol Biol. 2009;16:130–7.

40. Hall MP, Nagel RJ, Fagg WS, Shiue L, Cline MS, Perriman RJ, et al. Quaking and PTB control overlapping splicing regulatory networks during muscle cell differentiation. RNA. 2013;19:627–38.

41. Amir-Ahmady B, Boutz PL, Markovtsov V, Phillips ML, Black DL. Exon repression by polypyrimidine tract binding protein. RNA. 2005;11:699–716.

42. Collado-Torres L, Nellore A, Kammers K, Ellis SE, Taub MA, Hansen KD, et al. Reproducible RNA-seq analysis using recount2. Nature Biotechnology. 2017;35:319–21.

43. Melé M, Ferreira PG, Reverter F, DeLuca DS, Monlong J, Sammeth M, et al. Human genomics. The human transcriptome across tissues and individuals. Science. 2015;348:660–5.

44. Hentze MW, Castello A, Schwarzl T, Preiss T. A brave new world of RNA-binding proteins. Nature Reviews Molecular Cell Biology. 2018;19:327.

45. Gerstberger S, Hafner M, Tuschl T. A census of human RNA-binding proteins. Nature Reviews Genetics. 2014;15:829.

46. Anczuków O, Krainer AR. Splicing-factor alterations in cancers. RNA. 2016;22:1285–301.

47. Haberman N, Huppertz I, Attig J, König J, Wang Z, Hauer C, et al. Insights into the design and interpretation of iCLIP experiments. Genome Biol. 2017;18:7.

48. Lee D, Zhang J, Liu J, Gerstein M. Epigenome-based splicing prediction using a recurrent neural network. PLoS Comput Biol. 2020;16:e1008006–e1008006.

49. Yearim A, Gelfman S, Shayevitch R, Melcer S, Glaich O, Mallm J-P, et al. HP1 is involved in regulating the global impact of DNA methylation on alternative splicing. Cell Rep. 2015;10:1122–34.

50. Ke S, Shang S, Kalachikov SM, Morozova I, Yu L, Russo JJ, et al. Quantitative evaluation of all hexamers as exonic splicing elements. Genome Res. 2011;21:1360–74.

51. Begg BE, Jens M, Wang PY, Minor CM, Burge CB. Concentration-dependent splicing is enabled by Rbfox motifs of intermediate affinity. Nature Structural & Molecular Biology. 2020;27:901–12.

52. Wu JI, Reed RB, Grabowski PJ, Artzt K. Function of quaking in myelination: regulation of alternative splicing. Proc Natl Acad Sci U S A. 2002;99:4233–8.

53. Chen F, Keleş S. SURF: integrative analysis of a compendium of RNA-seq and CLIP-seq datasets highlights complex governing of alternative transcriptional regulation by RNA-binding proteins. Genome Biol. 2020;21:139.

54. Carazo F, Gimeno M, Ferrer-Bonsoms JA, Rubio A. Integration of CLIP experiments of RNA-binding proteins: a novel approach to predict context-dependent splicing factors from transcriptomic data. BMC Genomics. 2019;20:521–521.

55. Havugimana PC, Hart GT, Nepusz T, Yang H, Turinsky AL, Li Z, et al. A census of human soluble protein complexes. Cell. 2012;150:1068–81.

56. Zhang Z, Pan Z, Ying Y, Xie Z, Adhikari S, Phillips J, et al. Deep-learning augmented RNA-seq analysis of transcript splicing. Nat Methods. 2019;16:307–10.

57. Krakau S, Richard H, Marsico A. PureCLIP: capturing target-specific protein-RNA interaction footprints from single-nucleotide CLIP-seq data. Genome Biol. 2017;18:240.

58. Smith T, Heger A, Sudbery I. UMI-tools: modeling sequencing errors in Unique Molecular Identifiers to improve quantification accuracy. Genome Res. 2017;27:491–9.

59. Quinlan AR, Hall IM. BEDTools: a flexible suite of utilities for comparing genomic features. Bioinformatics. 2010;26:841–2.

60. Dale RK, Pedersen BS, Quinlan AR. Pybedtools: a flexible Python library for manipulating genomic datasets and annotations. Bioinformatics. 2011;27:3423–4.

61. Cock PJA, Antao T, Chang JT, Chapman BA, Cox CJ, Dalke A, et al. Biopython: freely available Python tools for computational molecular biology and bioinformatics. Bioinformatics. 2009;25:1422–3.

62. Bailey TL. DREME: motif discovery in transcription factor ChIP-seq data. Bioinformatics. 2011;27:1653–9.

63. Heller D, Krestel R, Ohler U, Vingron M, Marsico A. ssHMM: extracting intuitive sequence structure motifs from high-throughput. Nucleic Acids Res. 2017;45:11004–18.

64. Buschmann T. DNABarcodes: an R package for the systematic construction of DNA sample tags. Bioinformatics. 2017;33:920–2.

65. Gibson DG, Young L, Chuang R-Y, Venter JC, Hutchison CA 3rd, Smith HO. Enzymatic assembly of DNA molecules up to several hundred kilobases. Nat Methods. 2009;6:343–5.

66. Koblan LW, Doman JL, Wilson C, Levy JM, Tay T, Newby GA, et al. Improving cytidine and adenine base editors by expression optimization and ancestral reconstruction. Nat Biotechnol. 2018;36:843–6.

67. Kim D, Paggi JM, Park C, Bennett C, Salzberg SL. Graph-based genome alignment and genotyping with HISAT2 and HISAT-genotype. Nat Biotechnol. 2019;37:907–15.

68. Li H, Handsaker B, Wysoker A, Fennell T, Ruan J, Homer N, et al. The Sequence Alignment/Map format and SAMtools. Bioinformatics. 2009;25:2078–9.

69. Li H. A statistical framework for SNP calling, mutation discovery, association mapping and population genetical parameter estimation from sequencing data. Bioinformatics. 2011;27:2987–93.

70. Hwang G-H, Park J, Lim K, Kim S, Yu J, Yu E, et al. Web-based design and analysis tools for CRISPR base editing. BMC Bioinformatics. 2018;19:542.

71. Rausch T, Fritz MH-Y, Untergasser A, Benes V. Tracy: basecalling, alignment, assembly and deconvolution of sanger chromatogram trace files. BMC Genomics. 2020;21:230.

72. Lopez-Delisle L, Rabbani L, Wolff J, Bhardwaj V, Backofen R, Grüning B, et al. pyGenomeTracks: reproducible plots for multivariate genomic datasets. Bioinformatics. 2020. doi:10.1093/bioinformatics/btaa692.

73. Colaprico A, Silva TC, Olsen C, Garofano L, Cava C, Garolini D, et al. TCGAbiolinks: an R/Bioconductor package for integrative analysis of TCGA data. Nucleic Acids Research. 2016;44:e71–e71.

74. Friedman JH, Hastie T, Tibshirani R. Regularization Paths for Generalized Linear Models via Coordinate Descent. Journal of Statistical Software; Vol 1, Issue 1 (2010). 2010. https://www.jstatsoft.org/v033/i01.

75. Ashburner M, Ball CA, Blake JA, Botstein D, Butler H, Cherry JM, et al. Gene ontology: tool for the unification of biology. The Gene Ontology Consortium. Nat Genet. 2000;25:25–9.

76. Carbon S, Ireland A, Mungall CJ, Shu S, Marshall B, Lewis S. AmiGO: online access to ontology and annotation data. Bioinformatics. 2009;25:288–9.

77. The Gene Ontology Resource: 20 years and still GOing strong. Nucleic Acids Res. 2019;47:D330–8.

